# A Multiplex Protein Microarray Platform for Preclinical Tissue Binding and Cross-Reactivity of Therapeutic Antibodies

**DOI:** 10.64898/2026.10.11.757820

**Authors:** Susumu Matsukura, Naoki Sawada, Munehiro Watanabe, Yuya Tajima, Ibuki Yabuta, Gaku Morisawa, Yoshifumi Kawamura, Hideaki Nanamiya, Jun-ichi Imai, Motoki Takagi

**Author notes:** Correspondence: Motoki Takagi.

## Abstract

The NexAb multiplex protein microarray platform, comprising a 7,015-spot tissue lysate array and antigen-oriented protein arrays, was developed for preclinical tissue-binding and cross-reactivity assessment of therapeutic antibodies. Four anti-HER2 antibodies, trastuzumab, pertuzumab, trastuzumab emtansine (T-DM1), and trastuzumab deruxtecan (T-DXd), were profiled with integrated ERBB2 mRNA data. On antigen-oriented arrays, cell-synthesized HER2 bound trastuzumab and pertuzumab strongly and epitope-dependently, whereas wheat-germ HER2 showed no binding. On the tissue lysate array, trastuzumab and T-DM1 bound HER2-positive cancer cell lines and tumor tissues with nearly identical profiles (correlation coefficient r = 0.991), and T-DXd bound ERBB2-low cancer spots and 38.8% of normal human tissue spots, a profile not conserved in cynomolgus monkeys. Trastuzumab, T-DM1, and pertuzumab showed no cross-reactivity with cynomolgus monkey tissues, whereas T-DXd bound 3.1% of monkey tissue spots. NexAb enables quantitative cross-reactivity profiling relevant to preclinical safety evaluation.

---

Human epidermal growth factor receptor 2 (HER2; ERBB2/neu) is a cell-membrane-bound receptor tyrosine kinase and a representative therapeutic target for which gene amplification and protein overexpression are observed in multiple cancer types, including breast cancer. Anti-HER2 antibody therapy was established as the first molecularly targeted treatment for HER2-positive metastatic breast cancer through trastuzumab (Herceptin), a humanized monoclonal antibody that recognizes the extracellular domain IV of HER2 (Slamon et al. 2001). Subsequently, pertuzumab (Perjeta), which inhibits HER2 dimerization, prolonged overall survival in first-line treatment as part of a three-drug combination therapy with trastuzumab and docetaxel (Baselga et al. 2012). The antibody–drug conjugate (ADC) trastuzumab emtansine (T-DM1, Kadcyla) improved outcomes in previously treated HER2-positive breast cancer (Verma et al. 2012). Furthermore, trastuzumab deruxtecan (T-DXd, Enhertu), a next-generation ADC, showed superior progression-free and overall survival compared with T-DM1 (Cortés et al. 2024) and is now widely used as the new standard of care for HER2-positive breast cancer. All four of these agents are approved drugs, and their indications are expanding to include multiple cancer types beyond breast cancer. On the other hand, characteristic adverse events reported for these drugs include cardiotoxicity for trastuzumab-based agents (Nemeth et al. 2017; Mohan et al.,2018) and interstitial lung disease (ILD) for T-DXd (Modi et al.,2025); HER2 expression in normal tissues, including the myocardium, has been suggested to underlie cardiotoxicity (Nemeth et al. 2017).

An ADC is a complex in which a cytotoxic payload is linked to an antibody—which is specific to a target antigen on the surface of tumor cells—via a linker; it is designed to separate therapeutic efficacy from toxicity through antigen-dependent drug delivery (Panowski et al. 2014). The majority of conjugations utilize lysine residues or cysteine residues derived from interchain disulfides, and site-specific conjugation, which controls the conjugation site, has been reported to improve the therapeutic index (Junutula et al. 2008; Schumacher et al. 2016; Zhou et al. 2021). However, systematic findings regarding whether the act of conjugating a drug to an antibody itself alters antigen-binding affinity are limited. A study using model ADCs reported that differences in linker and conjugation chemistry did not result in significant changes in antigen-binding affinity (Acchione et al. 2012), whereas the conjugation site has been demonstrated to modulate the in vivo stability and therapeutic activity of ADCs (Shen et al. 2012); therefore, changes in binding characteristics associated with drug conjugation must be verified for each individual ADC. Since T-DM1 and T-DXd are based on the same trastuzumab backbone but differ in their linker, payload, and conjugation schemes, comparing their tissue-binding profiles is critical for evaluating the impact of conjugation on antigen recognition.

In fact, T-DXd is effective even in HER2-low breast cancer—previously classified as HER2-negative—indicating that detailed binding-profile analysis can guide the expansion of indications (Modi et al. 2025). In the development of antibody therapeutics, comprehensively understanding which cancer tissues a target antibody actually binds to. Furthermore, nonclinical safety evaluations of therapeutic antibodies require tissue cross-reactivity (TCR) testing based on the ICH S6(R1) guideline (ICH 2011; Fujii et al. 2017), and binding profiles in tissue panels from humans and relevant animal species are utilized in the selection of animal species for toxicity testing and in safety evaluations (Lynch et al. 2009). Conventional TCR studies have relied primarily on immunohistochemical evaluation of tissue sections, and few methods allow quantitative comparison of multiple antibodies across many tissue-derived samples under identical conditions.

Reverse-phase protein array (RPPA) technology enables quantitative comparison of antibody binding by spotting numerous tissue and cell lysates on a single substrate and has proven useful in clinical and basic research (Akbani 2014). We have also developed forward-phase, antigen-oriented protein arrays: an 18K protein array carrying proteins synthesized from human cDNAs in a wheat-germ cell-free expression system (18,429 spots) and a 5.3K protein array carrying proteins synthesized in cells (5,376 spots). These arrays have been applied to the identification of novel autoantibodies in patients with IPEX syndrome (Hoshino et al. 2019), the evaluation of vascular endothelial growth factor-binding peptides (Michigami et al., 2021), the profiling of salivary IgA antibodies against human proteins in patients with COVID-19 (Hikichi & Kunieda 2026), and the identification of autoantibodies specific for autoimmune hepatitis (Abe et al. 2025). In this study, we constructed the NexAb platform—a multiplex protein microarray platform comprising these antigen-oriented arrays and a multi-sample tissue lysate array constructed in the RPPA format, which carries protein extracts derived from human tumor tissues, normal tissues, cancer cell lines, and normal cynomolgus monkey tissues and enables integrated analysis with ERBB2 gene expression data. Prior to tissue profiling, the antigen recognition of the antibodies was validated on the antigen-oriented arrays, on which HER2 proteins synthesized in different expression systems were spotted, and the influence of direct fluorescent labeling of the antibody on the binding profiles was examined. We then systematically evaluated the tissue-binding profiles and tissue cross-reactivity of four anti-HER2 monoclonal antibodies on the NexAb platform.

## Materials and methods

### Antibodies

Trastuzumab (Herceptin; Roche, Basel, Switzerland; lot no. H4569), pertuzumab (Perjeta; Roche, Basel, Switzerland; lot no. H0108B06), trastuzumab emtansine (T-DM1, Kadcyla; Roche, Basel, Switzerland; lot no. N1019B10), and trastuzumab deruxtecan (T-DXd, Enhertu; Daiichi Sankyo, Tokyo, Japan; lot no. WJA0002) were used in this study.

### Fluorescent labeling of antibodies

For direct labeling, trastuzumab and pertuzumab were labeled with Alexa Fluor 647 using the Alexa Fluor 647 antibody labeling kit (Thermo Fisher Scientific, Waltham, MA, USA; cat. no. A20186).

### Protein microarrays

All microarray experiments in this study were performed using a protein microarray platform originally developed at Fukushima Medical University. Three microarrays were used in this study: the tissue lysate array (reverse-phase format), an 18K forward-phase antigen array carrying human proteins synthesized from human cDNAs in a wheat-germ cell-free expression system, and a 5.3K forward-phase antigen array carrying human proteins synthesized in cells (Supplementary Tables S1 and S2). ERBB2 mRNA expression data obtained by an independent transcriptome analysis were available for 5,755 spots (82.0% of the array) on the lysate array.

All microarrays were processed under the same conditions using a protein microarray incubation cassette (P-DE1; Fukushima Protein Factory, Inc., Fukushima, Japan; cat. no. HC4-01-DE). The arrays were washed with 100% ethanol, the final wash solution for protein microarrays (Fukushima Protein Factory, Inc.; cat. no. PA1210), and Solution A (Fukushima Protein Factory, Inc.; cat. no. PA1010), and then blocked with Blocking One (Nacalai Tesque, Inc., Kyoto, Japan; cat. no. 03953-95) for 1 h at 26°C with rocking. After washing with Solution A, the arrays were incubated for 17 h at 37°C with rocking in the primary antibody dilution buffer (Fukushima Protein Factory, Inc.; cat. no. PA1302) containing a fluorescently labeled antibody sample (1 µg/mL) and Goat Reference Antibody Mixture II (Fukushima Protein Factory, Inc.; cat. no. PA5011; 1:2,000), followed by washing with Solution A. The arrays were then incubated for 1 h at 26°C with rocking in the secondary antibody dilution buffer (Fukushima Protein Factory, Inc.; cat. no. PA1402) containing Cy3-conjugated anti-Goat IgG (Jackson ImmunoResearch Laboratories, Inc., West Grove, PA, USA; cat. no. 805-165-180; 1:500), washed with Solution A, Solution B (Fukushima Protein Factory, Inc.; cat. no. PA1110), and the final wash solution (cat. no. PA1210), and dried by centrifugation at 250 × *g* for 1 min. When unlabeled antibodies were evaluated on the antigen-oriented protein arrays, Alexa Fluor 647-labeled anti-human IgG was used as the secondary antibody for the detection of bound human IgG, together with Cy3-conjugated anti-Goat IgG for reference detection.

The arrays were scanned using a GenePix 4000B scanner (Molecular Devices, San Jose, CA, USA). The fluorescence intensity ratios of Alexa Fluor 647 (635 nm) to Cy3 (532 nm) were calculated as the median of ratios and expressed as relative log_2_ values, enabling comparison among the samples on the same array. Because protein microarray measurements provide relative quantification, binding levels were interpreted as relative signal intensities among the samples. For reverse-phase microarrays, any Alexa Fluor 647 measurement (raw fluorescence intensity) where the difference between the array used to evaluate the antibody and the negative control array is less than 20 is replaced with 0.

### Ethics approval and consent to participate

All experiments involving materials derived from humans were performed in accordance with the Declaration of Helsinki and were approved in advance by the Ethics Committee of the Fukushima Medical University (Fukushima, Japan; approval nos. 1953 and 2192; approval dates March 18, 2020, and May 26, 2016, respectively). The participants provided written informed consent.

### Use of Artificial Intelligence Tools

Large language model support was used during manuscript preparation through Genspark (MainFunc Inc. / Genspark Inc.; https://www.genspark.ai/) to assist with outlining and drafting selected text passages. All generated output was reviewed, revised, and verified by the authors against original sources. The authors take full responsibility for the final content of the manuscript.

## Results

### NexAb: two-step antibody profiling—from antigen-oriented validation to tissue binding

The NexAb platform is a multiplex protein microarray platform designed to profile therapeutic antibodies in two consecutive steps (Figure 1). In the first step, antigen-oriented protein arrays—an 18K forward-phase array carrying 18,429 protein spots synthesized from human cDNAs in a wheat-germ cell-free expression system and a 5.3K array carrying 5,376 protein spots synthesized in cells—are used to confirm whether an antibody binds its target antigen and to assess its cross-reactivity with proteins other than the target (Figure 2A). In the second step, the antibody is applied to a multi-sample tissue lysate array—a RPPA in which numerous tissue and cell lysates are spotted—to determine which tissues the antibody actually binds (Figure 2B and C). Because all arrays are processed under identical conditions, the results of the two steps can be interpreted on a common scale of relative binding levels.

**Figure 1.**
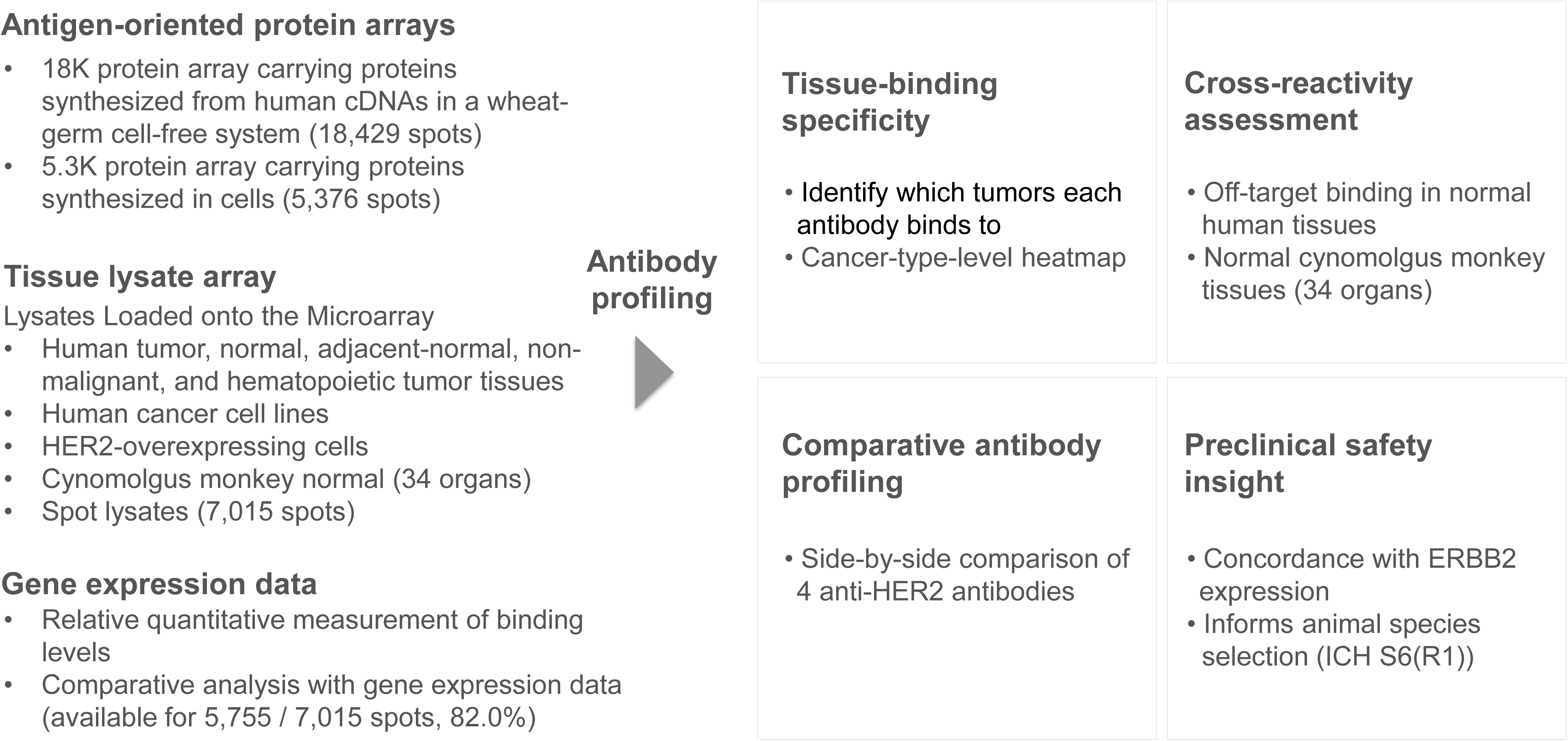
Overview of the NexAb platform. Antigen-oriented protein arrays validate antigen recognition, and the tissue lysate array profiles tissue binding; the platform yields four complementary read-outs—tissue-binding specificity, cross-reactivity assessment, comparative antibody profiling, and preclinical safety insight.

**Figure 2.**
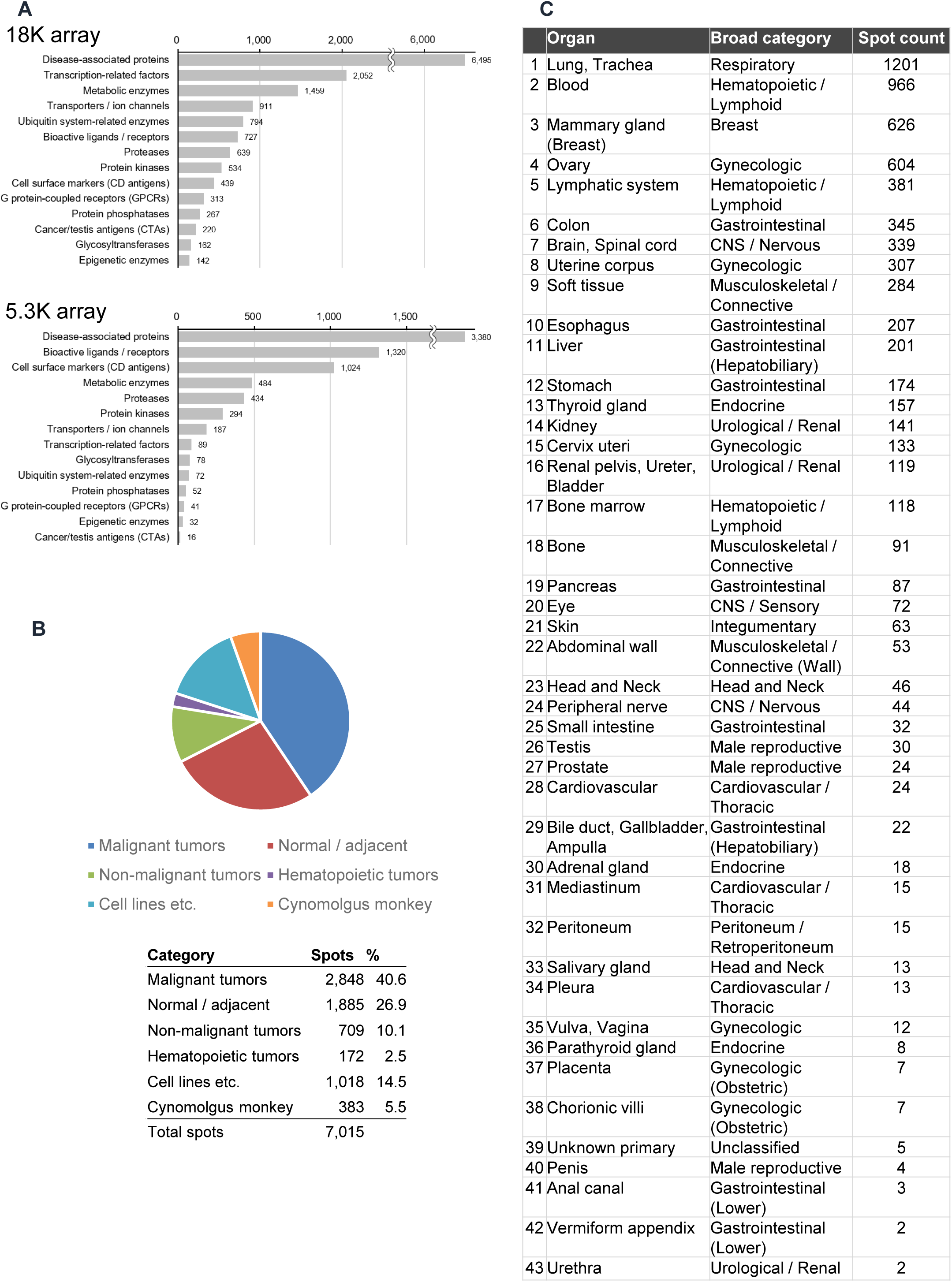
(A) Numbers of proteins loaded on the antigen-oriented protein arrays, grouped by functional category (upper panel, the 18K array; lower panel, the 5.3K array). The numbers shown in each category represent the number of genes. There are genes that appear in more than one category. (B) Composition of the arrayed samples: malignant tumors (2,848 spots; 40.6%), normal and tumor-adjacent tissues (1,885 spots; 26.9%), non-malignant tumors (709 spots; 10.1%), hematopoietic tumors (172 spots; 2.5%), cell lines and related specimens (1,018 spots; 14.5%), and normal cynomolgus monkey tissues (383 spots; 5.5%). (C) Number of spots per organ or tissue type (43 tissue types in total).

The antigen-oriented protein arrays are forward-phase arrays on which purified proteins are spotted and probed with the test antibody, so that binding to individual proteins can be assessed independently of tissue context. The proteins loaded on the two arrays are classified into 14 functional categories, among which disease-associated proteins form the largest category, followed by transcription-related factors, metabolic enzymes, transporters and ion channels, and bioactive ligands and receptors (Figure 2A). This category-level composition defines the antigen space that is interrogated for target binding and for cross-reactivity with non-target proteins before tissue profiling.

The tissue lysate array contains 7,015 spots of lysates prepared from human clinical specimens, human cell lines, and normal cynomolgus monkey tissues. Since protein microarray measurements provide relative quantification, antibody binding levels were determined and compared as relative signal intensities among the arrayed samples. The composition of the array is shown in Figure 2B: malignant tumors (2,848 spots; 40.6%), normal and tumor-adjacent tissues (1,885 spots; 26.9%), non-malignant tumors (709 spots; 10.1%), hematopoietic tumors (172 spots; 2.5%), cell lines and related specimens (1,018 spots; 14.5%), and normal cynomolgus monkey tissues (383 spots; 5.5%). The array covers 43 organ and tissue types (Figure 2C), with the largest numbers of spots for lung/trachea (1,201), blood (966), mammary gland (626), ovary (604), and the lymphatic system (381). Using this array, the binding profiles of the four anti-HER2 antibodies were obtained as relative binding levels for each arrayed sample, enabling side-by-side comparison among antibodies under identical conditions. *ERBB2* mRNA expression data obtained from an independent transcriptome analysis were integrated with the binding signals for 5,755 spots (82.0% of the array), enabling direct, spot-level comparison between gene expression and relative antibody binding. Collectively, the NexAb platform provides integrated insights into therapeutic antibodies with respect to tissue-binding specificity, cross-reactivity assessment, comparative antibody profiling, and preclinical safety insight (Figure 1).

### Validation of antigen binding and the effect of fluorescent antibody labeling on antigen-oriented protein arrays

To validate antigen recognition by the antibodies at the protein level, we first examined binding using an 18K forward-phase protein array carrying 18,429 protein spots synthesized from human cDNAs in a wheat-germ cell-free expression system, which included wild-type ERBB2 (four independent clones) and 13 ERBB2 mutant constructs (Supplementary Table S1). Unlabeled trastuzumab and unlabeled pertuzumab, detected with a fluorescent secondary antibody, showed no detectable binding to any of the 17 ERBB2 spots (all signals, 0.0), and Alexa Fluor 647-labeled trastuzumab gave only a marginal signal (maximum signal, 1.15, for a single construct) (Figure 3A). Thus, HER2 proteins synthesized in the wheat-germ cell-free system were not recognized by either antibody, presumably because of the lack of post-translational modification and/or incomplete folding of the extracellular domain.

**Figure 3.**
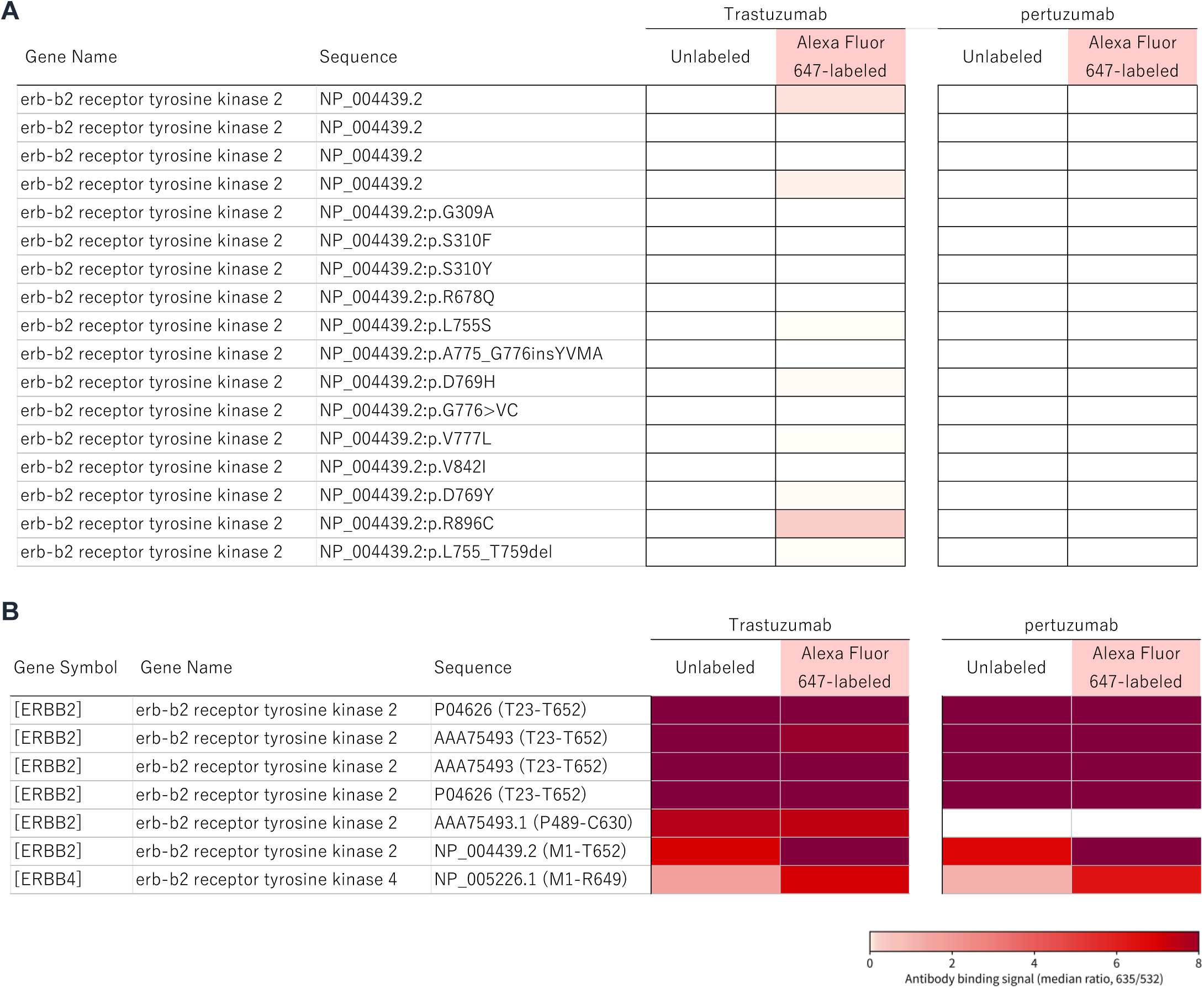
Validation of antigen binding on the antigen-oriented protein arrays. (A) Binding of unlabeled and Alexa Fluor 647-labeled trastuzumab and pertuzumab to the 17 ERBB2 spots on the 18K array. (B) Binding of trastuzumab and pertuzumab to the seven ERBB2 constructs on the 5.3K array, including the construct lacking the domain II epitope region (P489-C630) and the domain I construct (M1-C195).

We next used a 5.3K protein array carrying 5,376 proteins synthesized in cells, which included seven ERBB2 constructs (Supplementary Table S1). In marked contrast to the 18K array, both trastuzumab and pertuzumab bound strongly to the ERBB2 constructs covering the extracellular domain (signals up to 12.4 and 11.9, respectively) (Figure 3B). Pertuzumab did not bind to a construct lacking its domain II epitope region (P489-C630; trastuzumab 8.5 vs pertuzumab 0.0), confirming that antigen recognition was epitope-dependent. Off-target reactivity was limited on both arrays and overlapped substantially between the two antibodies (unlabeled antibodies: 3 of 18,412 proteins each on the 18K array; 12 of 5,368 and 5 of 5,368 proteins on the 5.3K array); ERBB4, a member of the same receptor family, was among the shared off-target proteins (Supplementary Table S1).

To evaluate the influence of direct fluorescent labeling of the antibody on profiling, we compared the protein-binding profiles obtained with the Alexa Fluor 647-labeled antibodies with those obtained with the unlabeled antibodies (detected with a labeled secondary antibody) on the 5.3K array. The profiles were highly concordant (r = 0.852 for trastuzumab and r = 0.899 for pertuzumab, Supplementary Table S1), although slightly more off-target proteins with weak positive signals were detected with the labeled antibodies on the 18K array (112 for trastuzumab and 29 for pertuzumab). These results validate the antigen specificity of the antibodies and indicate that direct fluorescent labeling has little effect on protein-binding profiles, supporting the reliability of the antibody profiling obtained on the NexAb platform.

### Profiling of the four anti-HER2 antibodies in cancer tissues and cell lines

Notably, binding to HER2-overexpressing cells, which are otherwise identical to the parental cells except for ERBB2 expression, demonstrates that the detected signals are target-dependent. This difference likely reflects the weaker binding of pertuzumab to HER2 relative to trastuzumab and T-DM1: on a lysate array, where arrayed proteins may be partially denatured and antigen density per spot is limited, an antibody with lower intrinsic affinity for its epitope is more likely to fall below the detection threshold. The overall antibody profiling data for the four anti-HER2 antibodies across the NexAb platform are presented in Supplementary Table S2. Trastuzumab, T-DM1, and T-DXd bound to HER2-positive cancer cell lines and to cells with forced HER2 expression (Figure 4A), confirming the validity of the NexAb platform for assessing antibody binding. On the tissue lysate array, in contrast, pertuzumab did not bind to the HER2-forced-expression cell lines, although it did bind to several HER2-positive cancer cell lines (Figure 4A). In addition, among cells expressing wild-type ERBB2 or ERBB2 variants, pertuzumab showed no detectable binding to cells expressing ERBB2 S310F, ERBB2 S310Y, or ERBB2 G309A—all of which lie within the domain II epitope recognized by pertuzumab—whereas trastuzumab, T-DM1, and T-DXd maintained binding to these variants.

**Figure 4.**
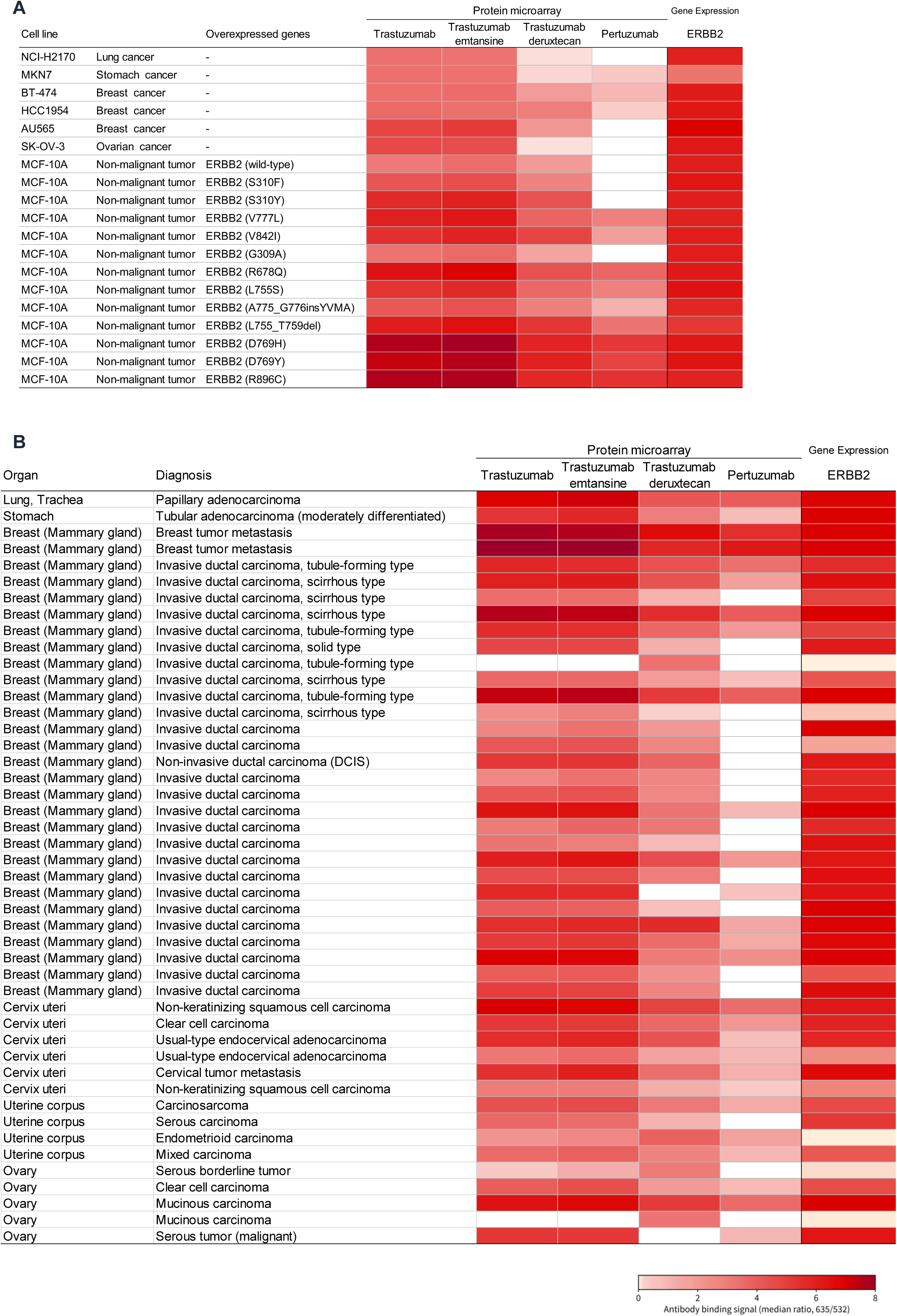
Tissue-binding profiles of the four anti-HER2 antibodies. (A) Binding to HER2-positive cancer cell lines, HER2-overexpressing cells, and ERBB2 variant-expressing cells. (B) Heatmap of the binding signals for the 46 cancer tissue spots with a signal of 3.0 or higher for at least one antibody, together with ERBB2 mRNA expression.

Among the malignant tumor tissue spots, those showing a binding signal of 3.0 or higher with at least one of the four antibodies were selected, yielding 46 cancer tissue spots for comparative profiling (Figure 4B). Of these, 38 (82.6%) showed high ERBB2 mRNA expression (ERBB2 ≥3). All of these were positive for trastuzumab (signal ≥1), 100% for T-DM1, 94.7% for T-DXd and 65.8% for pertuzumab. All four antibodies bound to HER2-positive tumor tissues, but their binding patterns differed. The binding profile of trastuzumab was nearly identical to that of T-DM1 (r = 0.991), consistent with the fact that both share a common antibody backbone and epitope. In contrast, the profile of T-DXd was clearly different from that of trastuzumab and T-DM1 (r = 0.460 and 0.479, respectively), showing more extensive binding that extended even to cancer tissues with low ERBB2 expression (ERBB2 < 3, n = 8). In these samples, T-DXd bound to all 8 spots (100%; median 2.97), whereas trastuzumab and T-DM1 bound 75.0% and pertuzumab bound 37.5%. Pertuzumab, which recognizes a different epitope (domain II) than trastuzumab and binds HER2 with lower intrinsic affinity, also exhibited a distinct profile, showing the lowest binding intensity among the four antibodies. Taken together, these results demonstrate that the NexAb platform can distinguish differences in binding profiles to cancer tissue among the four approved anti-HER2 drugs.

### Correlation between ERBB2 gene expression and antibody binding signals

To investigate the relationship between target gene expression and antibody binding, we integrated ERBB2 mRNA expression with antibody binding profiles at the spot level. In 5,755 tissue spots where ERBB2 expression data and binding data were paired, the binding signals for trastuzumab (r = 0.593) and T-DM1 (r = 0.597) showed a moderate positive correlation with ERBB2 mRNA expression, whereas for T-DXd (r = 0.343) and pertuzumab (r = 0.374), the correlation was weak (Figure 5). Consistent with this, binding signals increased as expression levels rose, but the extent of this increase varied by antibody.

**Figure 5.**
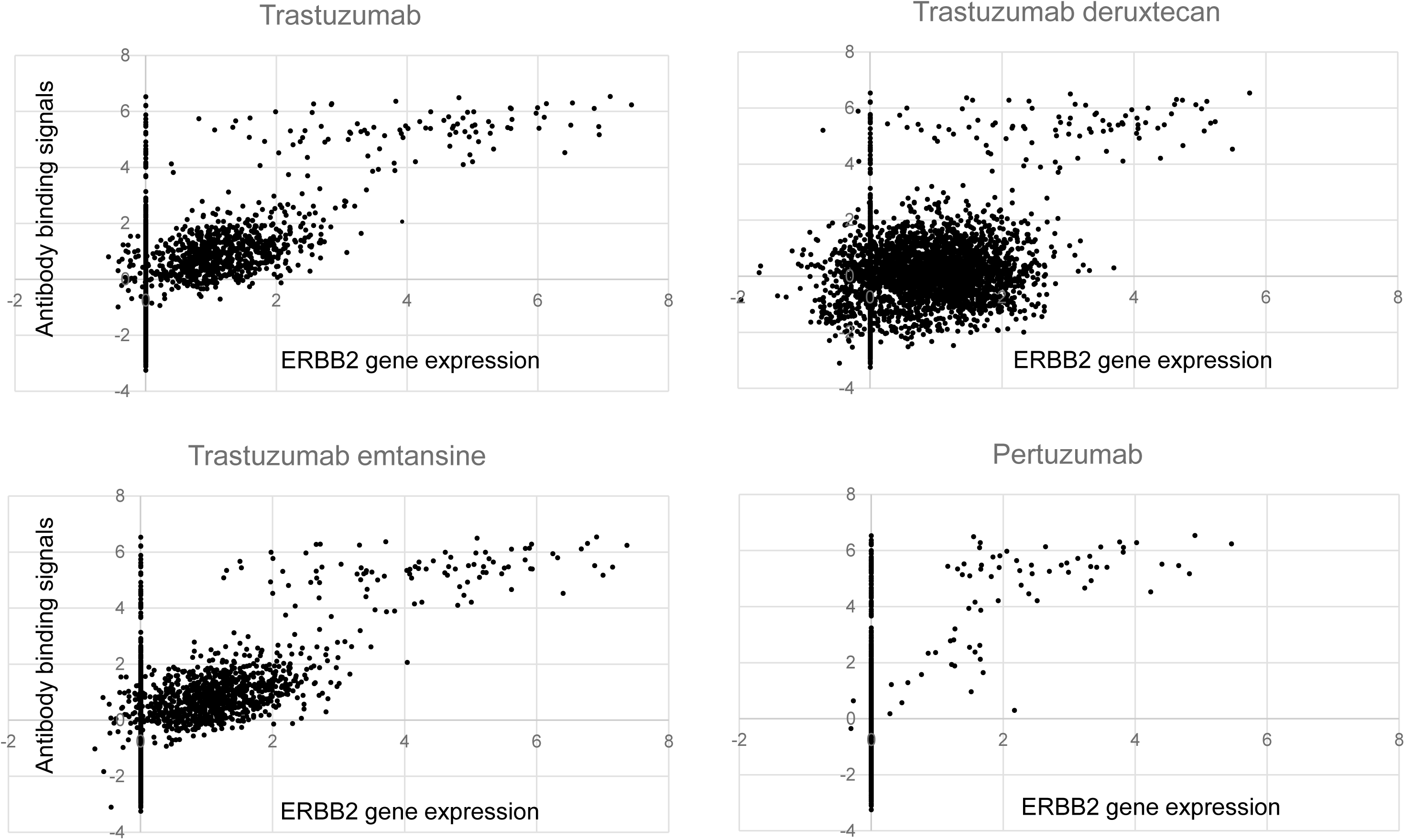
Relationship between ERBB2 gene expression and antibody binding signals. Correlation between ERBB2 mRNA expression and the binding signal for trastuzumab (r = 0.593), trastuzumab emtansine (r = 0.597), trastuzumab deruxtecan (r = 0.343), and pertuzumab (r = 0.374) across 5,755 tissue spots, together with the binding signals stratified by ERBB2 expression level, including the non-reactive spots among ERBB2-high spots (ERBB2 mRNA ≥ 4.0).

It is particularly noteworthy that high levels of ERBB2 mRNA expression do not fully predict antibody binding. Of the 115 spots with high ERBB2 expression (ERBB2 mRNA ≥ 4.0), 22 spots were non-reactive (binding signal = 0) for all four antibodies; the number of non-reactive spots was 25 for trastuzumab, 27 for T-DM1, 29 for T-DXd, and 68 for pertuzumab (Supplementary Table S2). The markedly higher number of non-reactive spots for pertuzumab is consistent with its lower intrinsic affinity for the domain II epitope of HER2, which may place its binding below the detection threshold in lysate spots with limited antigen density. These results suggest that factors other than the amount of target mRNA—such as antigen accessibility, tissue-dependent differences in protein expression, and the intrinsic affinity of the antibody—may influence antibody binding to tumor tissue.

### Antibody binding to normal tissues and cross-reactivity assessment

The cross-reactivity of the four anti-HER2 antibodies was evaluated by measuring their binding to normal human tissues (1,044 spots from 28 tissue samples; Table 1) and normal cynomolgus monkey organs (383 spots from 34 organs; Table 2), with spots exhibiting a binding signal of 1.0 or higher defined as positive. Pertuzumab showed no binding whatsoever to spots from normal human tissues (0 out of 1,044 spots, 0.0%). This lack of binding was similarly observed in monkey organs (0 out of 383 spots). These results demonstrate that pertuzumab exhibits virtually no cross-reactivity with normal tissues in either species.

In normal human tissues, trastuzumab and T-DM1 were positive at a limited number of sites: 52 out of 1,044 (5.0%) and 77 out of 1,044 (7.4%) respectively. The positive sites were concentrated in the lungs (18 and 24), stomach (15 and 19), colon (10 and 16), esophagus (5 and 7), and kidneys (3 and 7). In contrast, T-DXd was positive in 405 of 1,044 spots (38.8%), with binding distributed across 21 tissues. The most prominent sites were the lungs (238), the colon (32), the breast (31), the stomach (23), and the ovaries (20), followed by the esophagus (11), the thymus (10), the uterus (9), urothelial tissue (8), and the kidneys (8). In addition, T-DXd was positive in 2 of 3 biliary tract spots, whereas one positive spot each was detected in the bone, eye, mediastinum, penis, pleura, proctodeum, testis, and vulva/vagina. T-DXd therefore showed markedly broader cross-reactivity in normal human tissues than the other three antibodies.

In normal organs of cynomolgus monkeys, the binding profiles differed significantly from those in humans. While trastuzumab, T-DM1, and pertuzumab showed no positive binding at any of the 383 organ spots, T-DXd showed positive binding at only 12 of the 383 spots (3.1%), and its distribution was limited to four organs: the cardiovascular system (5 out of 24 spots), the eye (4 out of 28), soft tissue (2 out of 45), and the mammary gland (1 out of 4). Of particular note is that the organ-level cross-reactivity of T-DXd was not conserved between the two species. Tissues in which T-DXd binding is frequently observed in humans (lungs, colon, breast, stomach, and ovaries) showed no positive spots in the corresponding organs of cynomolgus monkeys. This discrepancy between human and cynomolgus monkey tissue-binding profiles highlights the importance of simultaneous evaluation on a single platform when selecting animal species and interpreting nonclinical safety findings.

## Discussion

In this study, four clinically approved anti-HER2 antibodies directed against the same antigen were profiled under identical conditions on a single multiplex platform, and their binding behavior differed substantially in both malignant and normal tissues. Because all arrays were processed identically and interpreted on a common relative scale, these differences can be attributed to the antibodies rather than to assay variation. The platform yielded four complementary read-outs—tissue-binding specificity, cross-reactivity assessment, comparative antibody profiling, and preclinical safety insight.

Several observations support the validity of the platform. Binding to HER2-overexpressing cells, which differ from their parental cells only in ERBB2 expression, indicates that the signals are target-dependent rather than nonspecific adsorption. The epitope-specific behavior of pertuzumab—no binding to ERBB2 (S310F), ERBB2 (S310Y), or ERBB2 (G309A), all within its domain II epitope, whereas the domain IV-directed antibodies retained binding—reproduced antibody behavior consistent with the known epitope architecture of HER2. Pertuzumab nevertheless behaved differently from the other three antibodies on the tissue lysate array: it bound several HER2-positive cancer cell lines and purified mammalian-cell HER2, but not HER2-forced-expression lysates. This discrepancy is considered to reflect the lower intrinsic affinity of pertuzumab for HER2 (Adams et al. 2006; Lua et al. 2015), which should make it more sensitive to partial denaturation of its conformational epitope during lysate preparation and to the limited antigen density within a single spot; this interpretation is not a proven mechanism, and direct comparison of HER2 protein amounts in the corresponding spots would be required to substantiate it.

The antigen-oriented arrays provided independent validation at the purified-protein level and illustrated a key methodological consideration. Recognition of HER2 depended on the expression system: wheat-germ cell-free HER2 was not recognized, whereas mammalian-cell HER2 was bound strongly and epitope-dependently, indicating that conformational epitopes require correct folding and post-translational modification and that the choice of synthesis system is critical. The high concordance between labeled and unlabeled antibodies, together with the limited and largely overlapping off-target reactivity on both arrays, further supports the specificity of these antibodies. In cancer tissues, the four antibodies shared the same overall direction of binding but were clearly distinguishable in detail. The near-identity of the trastuzumab and T-DM1 profiles is consistent with their shared backbone and epitope, whereas the distinct, more extensive profile of T-DXd, which extended to ERBB2-low tissues, suggests that its binding depends less strictly on high target expression. Pertuzumab, recognizing a different epitope with lower affinity, consistently showed the lowest intensity. The platform therefore resolves antibody-to-antibody differences sufficient to discriminate agents sharing the same target.

The comparison between ERBB2 mRNA expression and binding signals further illustrated the capacity of the platform to detect target-expression-independent binding. Across 5,755 human tissue spots, binding correlated moderately with ERBB2 expression for trastuzumab (r = 0.593) and T-DM1 (r = 0.597) but only weakly for T-DXd (r = 0.343) and pertuzumab (r = 0.374). Importantly, among 115 spots with high ERBB2 expression (ERBB2 mRNA ≥ 4.0), 22 were non-reactive to all four antibodies, and the number of non-reactive spots per antibody ranged from 25 (trastuzumab) to 68 (pertuzumab). Gene expression therefore does not fully predict antibody binding, and the discordance between transcript-level and protein-level target detection—which conventional expression-based target assessment cannot capture—was directly visualized on this platform.

Cross-reactivity assessment in normal tissues demonstrated both the discriminative power and the translational relevance of the platform. The broader normal-tissue binding of T-DXd, in contrast to the essentially absent binding of the other three antibodies, is consistent with the distinct clinical toxicity profile reported for T-DXd, including ILD, and provides a plausible mechanistic basis for such off-tumor toxicities. Equally notable was the species discordance: the human normal-tissue profile of T-DXd was not reproduced in cynomolgus monkey organs. Because the cynomolgus monkey is the standard species for toxicology studies of anti-HER2 antibodies, this discordance cautions against extrapolating human off-tumor binding from monkey data alone and supports simultaneous evaluation of both species on a single platform.

Several limitations should be acknowledged. The panel composition is uneven, with rare tissues represented by only one to three spots, so organ-level conclusions for such tissues should be interpreted cautiously. The array measures binding to lysate-immobilized proteins, and tissue architecture, cell-surface accessibility, and antigen internalization are not reproduced, which may cause false-negative or false-positive binding; lower-affinity antibodies are especially susceptible, as exemplified by pertuzumab, and validation against immunohistochemistry or ex vivo tissue-section binding is warranted. Off-target signals for certain constructs may reflect expression-system artifacts. Binding was assessed relative to ERBB2 mRNA, and proteomic quantification of HER2 protein in the same spots would clarify the transcript–binding discordance. Because the thresholds were empirical and only relative quantification is possible, binding levels are comparative values rather than absolute affinities. Finally, only four anti-HER2 antibodies were profiled, and extending the platform to other targets and formats would establish its generalizability.

In summary, NexAb enables simultaneous, comparative profiling of antibody binding across malignant tissues, normal human tissues, and normal cynomolgus monkey organs, integrates this information with target gene expression, and detects antibody-specific and species-specific binding patterns that are relevant to preclinical safety assessment. Together with antigen-level validation using antigen-oriented arrays, this platform may complement conventional tissue cross-reactivity studies and support species selection and safety interpretation in antibody drug development.

## Supplementary material

Supplementary Table S1. Antibody binding signals of unlabeled and Alexa Fluor 647-labeled trastuzumab and pertuzumab on the 18K and 5.3K antigen-oriented protein arrays.

Supplementary Table S2. Antibody binding signals of trastuzumab, trastuzumab emtansine, trastuzumab deruxtecan, and pertuzumab and ERBB2 gene expression for the tissue lysate array.

## Data availability

The data that support the findings of this study are available from the corresponding author upon reasonable request.

## Authors’ contributions

SM, YK, JI, and MT conceived and designed the study. SM, NS, MW, YT, IY, GM, YK, and HN performed the experiments and analyzed the data. MT wrote the manuscript. All authors read and approved the final manuscript.

## Acknowledgements

Not applicable.

